# Spatiotemporal regulation of LGN/NuMA and astral microtubules generate mirror symmetric spindle rotations

**DOI:** 10.64898/2026.08.20.745970

**Authors:** Janet Chenevert, Anne Rosfelter, Daniel Gonzalez-Suarez, Silvia Caballero-Mancebo, Vlad Costache, Paul Stolz, Lydia Besnardeau, Rémi Dumollard, Alex McDougall

## Abstract

The positioning of the mitotic spindle controls the size, content, and position of daughter cells within embryos and tissues. A major spindle positioning mechanism is “cortical pulling” whereby the membrane-bound complex LGN/NuMA/Dynein captures astral microtubules and pulls centrosomes toward the cell cortex. Cytoplasmic dynein and astral growth tend to counteract cortical pulling and position spindles at the cell center. It remains unclear how these opposing forces cooperate. Here we examine the ascidian embryo, where spindles of two germ line cells rotate toward one another causing divisions which are both mirror symmetric and unequal. We find that this spindle behavior is governed by transient enrichment of LGN and NuMA and enhanced cortical pulling at the shared cell contact. Inhibition of the LGN/NuMA complex disrupts spindle alignment, unequal cleavage, and mirror symmetry. Temporal analysis shows that cortical pulling force initates at anaphase when there is a sharp increase in astral microtubule length. These results point towards two phases of spindle positioning forces, with cytoplasmic pulling and cortical pulling operating sequentially during early and late mitosis.

## Introduction

The positioning of the mitotic spindle guides the orientation of cell division and contributes in many substantial ways to the morphogenesis of a developing organism (reviewed in Williams and Fuchs, 2013; McDougall et al., 2019; Lechler and Mapelli, 2021; Sallé and Minc, 2022). Spindle position determines whether cell components including developmental determinants are partitioned equally between daughter cells or asymmetrically to generate nonidentical daughters and increase cell type diversity. It also controls the sizes of daughter cells which in turn influence embryo shapes. Spindle position dictates the initial positions of newborn cells within embryos or tissues and which neighbors they will contact. Finally, the coordination of spindle positions among groups of cells allows the planar expansion of epithelia and alignment with overall embryonic polarity.

The position of the spindle is controlled by forces exerted on astral microtubules (MTs), based on pulling by motors anchored in the cortex or in the cytoplasm, or on pushing due to MT growth. Recent work suggests that a tug of war between centering and off-centering forces orchestrates spindle movements to achieve the position required for each specific type of cell division. Centration of asters and spindles is favored by dynein in the cytoplasm pulling on the longest MTs and by pushing against the cell surface (Kimura and Kimura, 2011; Tanimoto et al., 2016; Pierre et al., 2016; Wu et al., 2017; Sallé et al., 2019; Meaders and Burgess, 2020; Meaders et al., 2020; Rosfelter et al., 2024). In contrast, cortical pulling on astral MTs can override centration forces in order to orient the spindle during asymmetric cell divisions or shift the spindle off-center for unequal cell division (UCD) (Gotta et al., 2003; Labbé et al., 2004; Redemann et al., 2010; Williams and Fuchs, 2013; Poon et al., 2019; Sallé et al., 2019; Wu et al., 2024). Localized cortical pulling is achieved in many systems by the conserved proteins LGN and NuMA which activate dynein at specific sites on the cell cortex. The protein complex LGN/NuMA/dynein controls a myriad of spindle orientations in animal development including planar cell divisions in epithelia, apico-basal alignment for asymmetric stem cell divisions, and off-centering for embryonic UCDs (di Pietro et al., 2016; Nakajima, 2018; Kiyomitsu and Boerner, 2021; Tarannum et al., 2022; Zellag et al., 2025). Numerous studies have revealed diverse spatial cues which can localize LGN or NuMA in different cell contexts, but how cortical pulling is integrated with other opposing forces and temporally with cell cycle phases are not well understood.

In bilaterian animals many structures and organs display mirror symmetry, meaning they are divided into two halves that are exact reflections of one another across a central axis. In the developing neural tube of the zebrafish embryo, spindles of progenitor cells orient perpendicular to a central axis, then after cell division the daughter cells polarize in a mirror image fashion towards their shared cleavage furrow which becomes part of the central axis (Tawk et al., 2007; Buckley and Clarke, 2014). In the *C. elegans* embryo, the gonad and vulva develop with mirror symmetry when specific cells reverse their anterior-posterior polarity in response to Wnt signalling and divide in opposite orientations (Green et al., 2008; So et al., 2024). In the annelid *Platynereis dumerilii*, mirror image structures of the larval brain arise from clonal progeny of multiple pairs of bilateral founder cells (Vopalensky et al., 2019). These few examples reveal a diversity of cellular strategies to generate mirror symmetry in developing embryos, but little is known about the spindle position mechanisms involved.

One striking example of mirror symmetric cell division is found in ascidian embryos, which display an invariant and highly conserved bilateral symmetry starting from first cleavage (Sardet et al., 2007; Negishi and Nishida, 2017; McDougall et al., 2019). In particular, the spindles of the two germline cells rotate toward one another causing divisions which are both mirror symmetric and unequal. Here we examined the role of the cortical pulling complex in the early embryo of the ascidian *Phallusia mammillata* (*Phallusia* hereafter), and found that the mirror image spindle rotations in neighbor germline cells depend on a transient enrichment of LGN and NuMA and enhanced cortical pulling at the shared cell contact. Temporal analysis reveals that while spindle rotation and off-centering take place throughout mitosis, cortical pulling initiates at anaphase. Our results point towards two distinct phases of force generation during early and late mitosis, mediated by shortening then lengthening of astral MTs respectively. This combination of mechanisms may explain how spindle position is both flexible yet precise over multiple cell divisions leading to the invariant cleavage pattern conserved among all ascidian embryos. We propose that such temporal regulation coupled with spatially localized cortical pulling position the spindle for precise mirror image UCD of germ line cells.

## Results and discussion

### Mirror symmetric spindle rotations in the ascidian germ lineage

Ascidian embryos exhibit an invariant cleavage pattern conserved in all species examined thus far. The first cleavage furrow creates a central axis of bilateral symmetry called the midline (red line in Fig. 1 A and dashed white line in Fig. 1 B and C) (Sardet et al., 2007; Negishi and Nishida, 2017; McDougall et al., 2019). Cell divisions proceed with mirror image symmetry across this axis, establishing identical left and right sides of the embryo. A cortical domain rich in determinant RNAs, germ plasm and cortical endoplasmic reticulum known as the centrosome attracting body or CAB (green in Fig. 1 C) is bisected by the first division and partitioned into a pair of cells at the 8-cell stage (Nishikata et al., 1999; Prodon et al., 2010; McDougall et al., 2015, 2019). These two posterior cells (called “B4.1” cells) then commence a series of three successive UCDs which segregate the CAB into the two smallest daughters. After the final UCD, the CAB-containing cells cease to divide, become internalized, and establish the primordial germ cells of the post-metamorphosis embryo (Shirae-Kurabayashi et al., 2006; Ohta and Christiaen, 2024).

**Figure 1.**
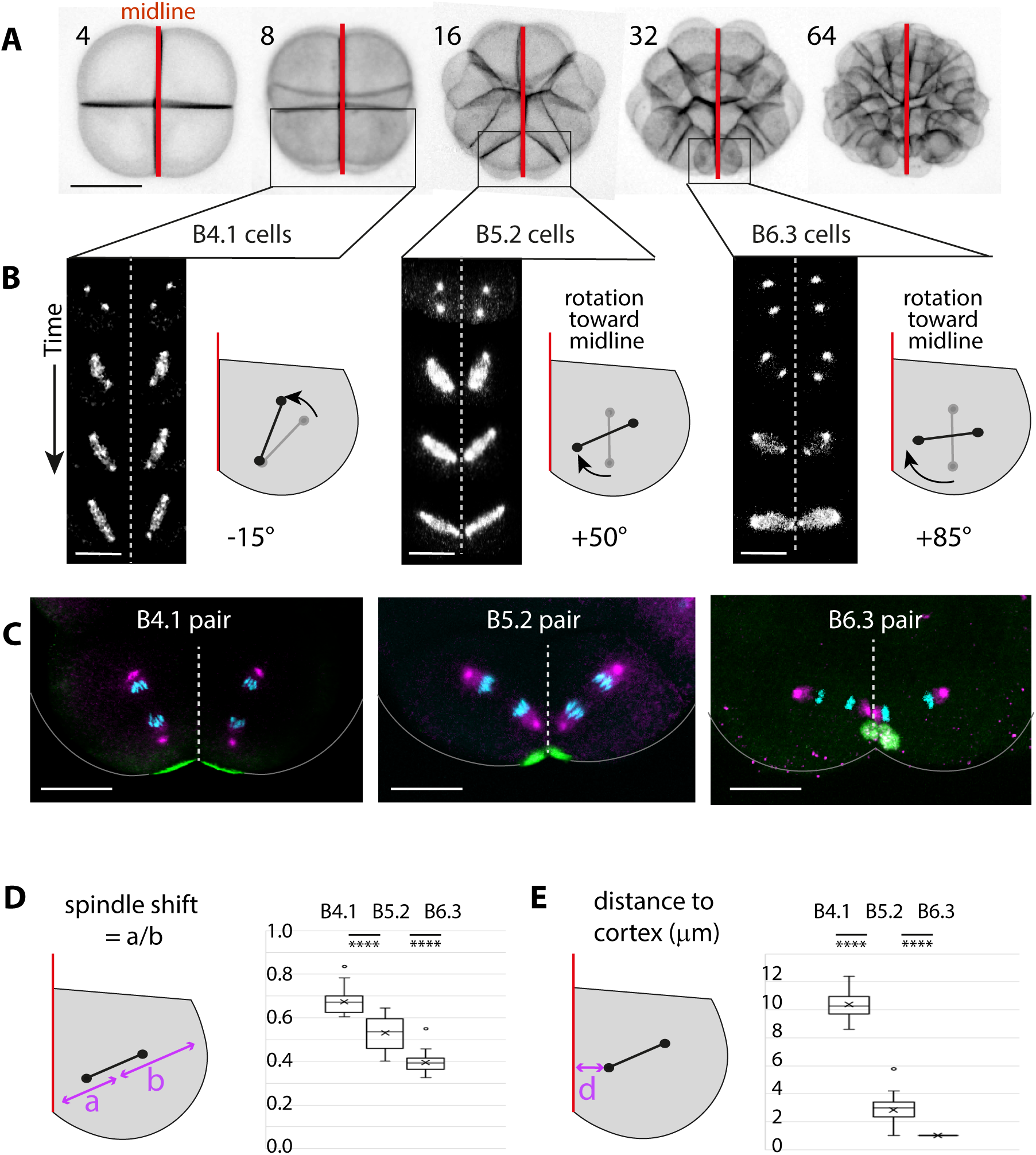
Mirror image spindle rotations toward the midline enhance unequal cell division. A) Projections of confocal Z-scans of a live *Phallusia* embryo labelled with Cell-Mask Green to label the plasma membrane and define cell contours. The number of cells in each embryo is indicated on top. The red line shows the midline axis of mirror symmetry. The two boxed cells each contain a CAB and undergo unequal cleavage. Scale bar = 60 microns. B) Images from timelapse videos of embryos expressing EB3-Venus (white) showing spindle behavior *in vivo*. Drawings represent the right cell of each pair showing spindle position at nuclear envelope breakdown (gray bars) and at anaphase (black bars). Spindles undergo rotation (curved arrows) toward the lateral midline in the B5.2 cells (second UCD) and B6.3 cells (third UCD), but not B4.1 cells (first UCD). The midline axis of mirror symmetry is indicated by a dotted white line in the image sequences or a red line in the drawings. Scale bar = 30 (left), 25 (middle), or 15 (right) microns. C) Images of fixed cells in anaphase labelled for spindle poles (anti-gamma tubulin, magenta), CAB domain (anti-aPKC, green) and DNA (Hoechst, cyan). The spindles point toward the midline during the second and third UCDs but not the first. Scale bar = 30 (left), 25 (middle), or 15 (right) microns, which is approximately anaphase spindle length for B4.1, B5.2, or B6.3 cells respectively. The midline axis of mirror symmetry is indicated by the white dotted line and apical cell contours are in gray. D) Degree of spindle shift at anaphase in each of the UCDs. The distance from the center of the spindle to the cell periphery along the spindle axis was measured in fixed samples (*n* = 20 cells for each stage). The ratio of the shortest distance (a) to the longest (b) gives the degree of spindle shift. One-way ANOVA test with Šídák’s multiple comparisons test (\*\*\*\**P*<0.0001). E) Distance (d) between midline cortex (in red) and closest centrosome in anaphase, measured in fixed samples immunolabelled with gamma tubulin antibody. *n* = 20 cells for each stage. One-way ANOVA test with Šídák’s multiple comparisons test (\*\*\*\**P*<0.0001).

Spindle movements in the germline cells were analyzed using live video microscopy and fixed immunofluorescence in the species *Phallusia mammillata*, whose eggs are transparent and translate exogenous RNAs thus favorable for live imaging of dynamic subcellular processes. During the first UCD of the series, the spindles in the two B4.1 cells form in the cell center then elongate toward the CAB while undergoing a slight change in angle with the midline plane (- 15°, Fig. 1 B-C, left). This spindle trajectory is in line with our previous study which showed that Kif2 localized in the CAB generates aster asymmetry and spindle shift (Costache et al., 2017). However during the second and third UCDs (B5.2 cells and B6.3 cells respectively), spindles form parallel to the midline then rotate toward the shared cell contact in a mirror image fashion (Fig. 1 B and C, center and right). With each sequential UCD the off-center spindle shift becomes more pronounced (Fig. 1 D) and the CAB-proximal centrosome moves closer to the midline cortex (Fig. 1 E), resulting in the two anaphase spindles pointing towards each other and not toward the CAB. We hypothesized that localized cortical pulling at this midline site adjacent to the CAB could cause these mirror image spindle rotations.

### Precise cortical pulling toward the midline cell contact between neighbor germ line cells

Sites of cortical pulling were directly visualized by the “membrane invagination” assay previously used in *C elegans* (Redemann et al., 2010; Berends et al., 2013, Fielmich et al., 2018) and *Phallusia* (Godard et al., 2021; Rosfelter et al., 2024). The rationale for this assay is that if the cell cortex is weakened by the addition of actin-depolymerizing drugs, cortically anchored dynein will bring attached plasma membrane toward the cell interior as it migrates toward the minus end of microtubules (Fig. 2 A). When 16-cell stage embryos fluorescently labelled for plasma membrane were treated with cytochalasin B and observed by video microscopy, prominent invaginations emerged from both sides of the midline directed toward the proximal centrosome in 94 % of analyzed B5.2 cells (Fig. 2 B-D green bars, arrows, Movie 1). Invaginations also formed, with less frequency, near the opposite spindle pole and on the apical side of the CAB (Fig. 2 C yellow bars, arrowheads in Fig. 2 D). These results show specific localized cortical pulling at the shared midline between the two CABs. To determine if this localized cortical pulling is dependent on LGN and NuMA, we examined their localization and function.

**Figure 2.**
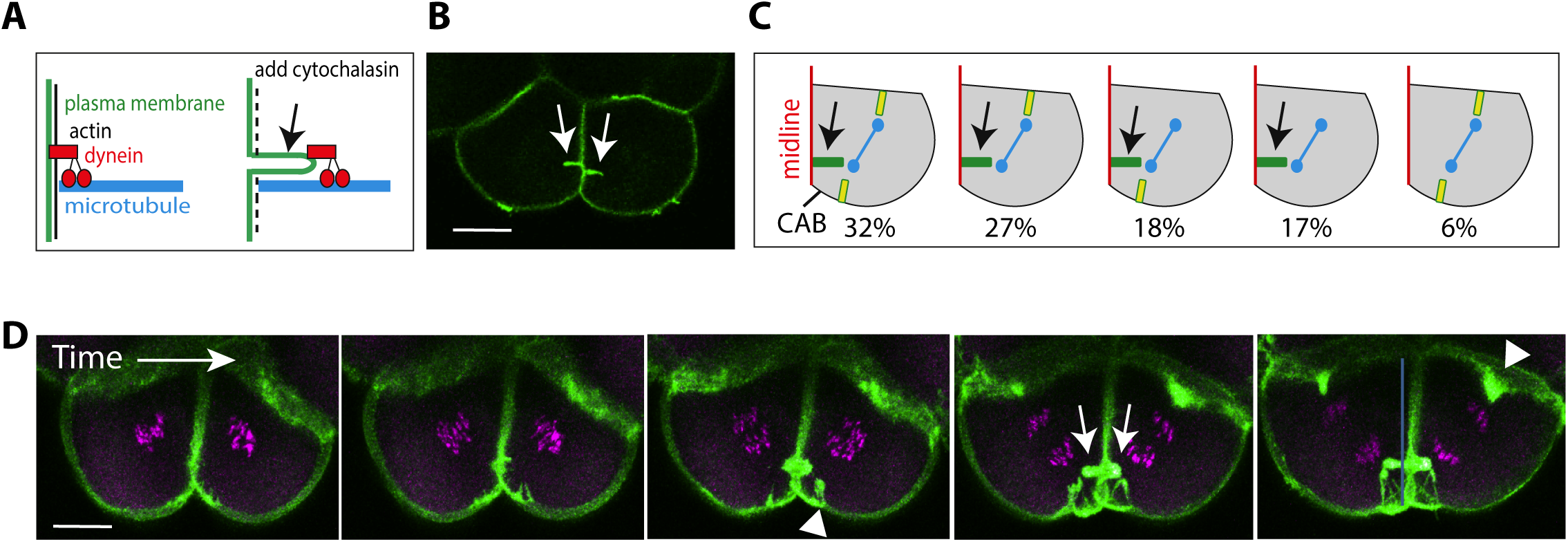
Localized cortical pulling at the midline between B5.2 cells. A) Schematic of the membrane invagination assay that reveals sites of cortical pulling. Arrow shows a membrane invagination which forms when the actin cortex is weakened and membrane-anchored dynein migrates toward the minus end of microtubules. B) A pair of B5.2 cells labelled with CellMask-Green and treated with cytochalasin D. Arrows indicate a major invagination in each cell emerging from the shared site of cell contact. Scale bar = 15 microns. C) Schematics of 5 patterns obtained in B5.2 cells treated with cytochalasin. One B5.2 cell of the pair is drawn, midline in red. Blue bar represents the spindle. Invaginations may appear at the midline (green bars), on the opposite pole (yellow bars, top right), or on the apical side (yellow bars on the bottom). Numbers indicate the frequencies of each observed pattern as percentages (*n* = 80 cells). D) Images from a timelapse film of a pair of B5.2 cells in an embryo treated with cytochalasin and expressing PH-GFP (green) to label plasma membrane and H2B-RFP (magenta) to label DNA. White arrows indicate the major midline invaginations (green bars in B and C), and arrowheads indicate minor invaginations (yellow bars in C). Time interval between images is 1 minute. Scale bar = 15 microns.

### LGN and NuMA accumulate at the B5.2 midline during mitosis

The distributions of LGN and NuMA proteins were evaluated using GFP constructs and specific antibodies. Ascidian LGN was identified by homology and cloned from a cDNA library of *Phallusia mammillata* or from the related ascidian species *Ciona intestinalis*. In live embryos expressing LGN tagged with Venus fluorescent protein at the N-terminus (Venus-LGN), LGN accumulated at the shared cell contact during mitosis in both B5.2 and B6.3 cell pairs (Fig. 3A). Endogenous LGN, recognized by an antibody made against full-length *Phallusia* LGN (Fig. S1 A), was similarly enriched at the shared midline in B5.2 and B6.3 cell pairs (Fig. 3 B, arrows). As LGN antibody was also observed to label the apical CAB domain, double staining with CAB markers (Patalano et al., 2006) was used to confirm that the midline signal identifies a region distinct from the CAB (Fig. 3 B; Movie 2). LGN antibody did not label cytoplasmic or microtubule structures, however in cytochalasin-treated embryos, endogenous LGN was found to accumulate on the centrosomes closest to the CAB (Fig. 3 C, arrows), consistent with traction between the LGN-rich midline cortex and the spindle pole. A similar observation was made in mammalian cells, where actin depolymerization caused cortical LGN to transit to the nearby centrosome (Zheng et al., 2013). Evaluation of endogenous LGN over the cell cycle in B5.2 cells revealed that midline enrichment increases in prometaphase and metaphase, peaks at anaphase and telophase, then is lost after cleavage (Fig. 3 D and E). Detailed spatial analysis at each timepoint showed that accumulation of LGN appears first near the apical face (CAB side), then increases inward along the midline (Fig. S1 B). During the subsequent UCD, LGN again localizes at the midline cell contact in B6.3 cells during mitosis and disappears after cleavage (Fig. 3 F, Fig. S1 C).

**Figure 3.**
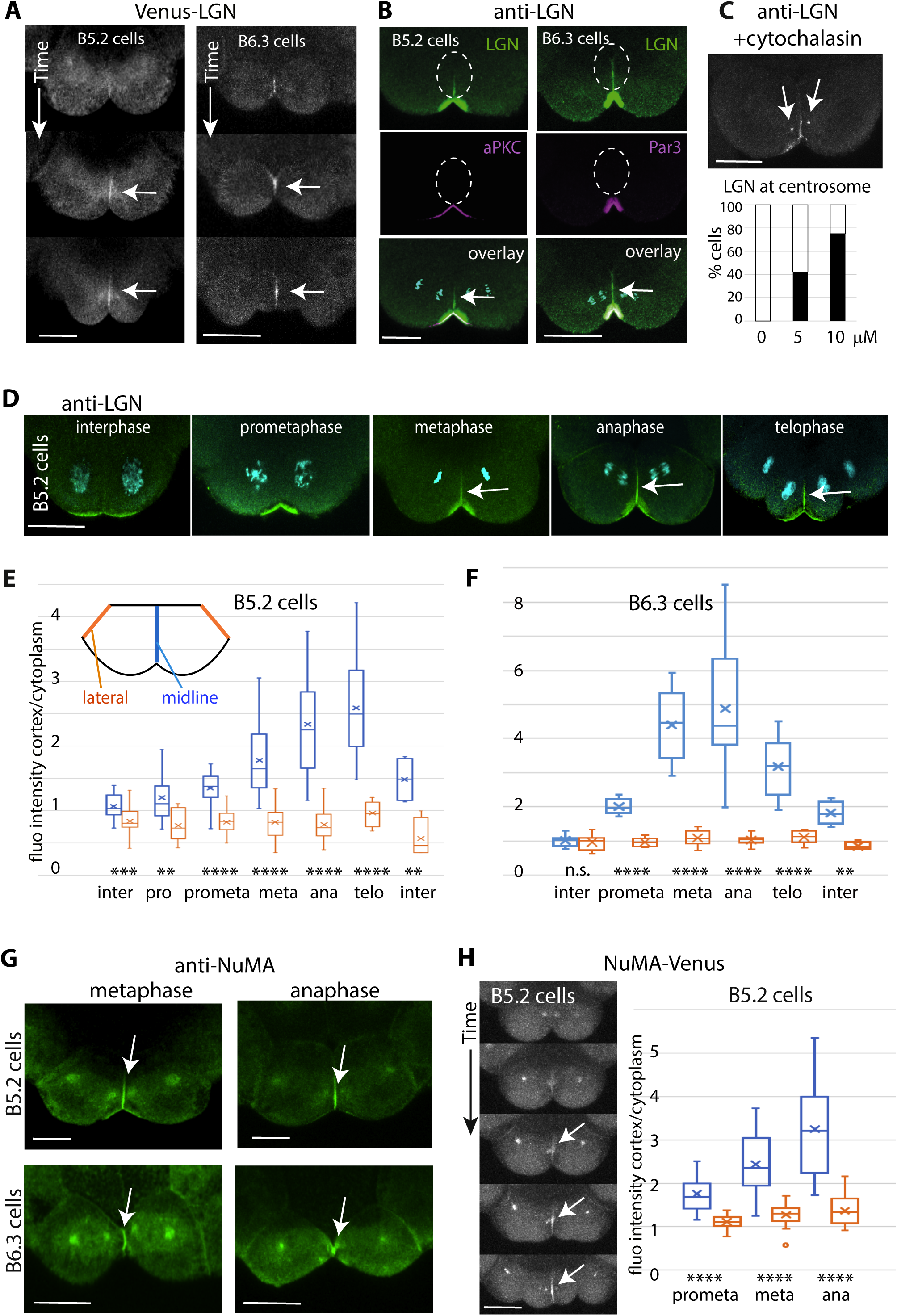
LGN and NuMA localize to the midline in B5.2 cells and B6.3 cells during mitosis. In all panels, scale bars = 20 microns. A) Images from timelapse movies of mitotic B5.2 cells (left) or B6.3 cells (right) expressing Venus-LGN (*Ciona* version). The time interval between images is 90 seconds. Horizontal arrows indicate LGN enrichment at the midline. B) Double labelling of a pair of B5.2 or B6.3 cells in anaphase with antibodies to LGN (green) and CAB markers aPKC or Par3 (pink). The DNA label (Hoechst, cyan) is added in the overlays (bottom images). White ovals indicate the position of the midline cell contact and arrows indicate enrichment of LGN at the midline. C) A pair of B5.2 cells from an embryo treated with cytochalasin then fixed in methanol and stained with anti-LGN (in white). Arrows indicate LGN signal at the centrosomes. The graph shows the frequency of cells with a labelled centrosome as a function of concentration of cytochalasin. *n* = 100 cells for 0 μM, 30 for 5 μM, and 46 for 10 μM cytochalasin. D) B5.2 cells fixed at the indicated phase of the cell cycle and stained with LGN antibody (green) and Hoechst (cyan). Arrows indicate LGN signal at the midline. E) Quantification of LGN accumulation in B5.2 cells labelled as in D, showing the ratio of fluorescence intensity at the midline (in blue) or lateral face (in orange) over cytoplasmic background. *n* = 21 cells for interphase, 13 for prophase, 25 for prometaphase, 40 for metaphase, 43 for anaphase, 13 for telophase, 4 for interphase after cleavage. Unpaired t test (\*\**P*=0.0016 for prophase; \*\**P*=0.0074 for interphase; \*\*\**P*=0.0004; \*\*\*\**P*<0.0001). F) Quantification of LGN accumulation in B6.3 cells labelled as in Fig. S1 C, showing the ratio of fluorescence intensity at the midline (in blue) or lateral face (in orange) over cytoplasmic background. *n* = 8 cells for interphase, 6 for prometaphase, 13 for metaphase, 24 for anaphase, 9 for telophase, 4 for interphase after cleavage. Unpaired t test (ns, non-significant, *P*=0.6; \*\**P*=0.0018; \*\*\*\**P*<0.0001). G) B5.2 cells (top row) or B6.3 cells (bottom row) from fixed embryos stained with anti-NuMA antibody (green). Arrows indicate NuMA signal at the midline. H) Images from a timelapse movie of B5.2 cells expressing NuMA-Venus (in white). The white arrow indicates NuMA signal at the midline. The time interval between images is 90 seconds. The graph shows quantification of NuMA-Venus accumulation during mitosis as the ratio of fluorescence intensity at the midline (in blue) or lateral face (in orange) over cytoplasmic background. Data from all B5.2 cells are grouped into cell cycle phases according to normalized time (see Methods). *n* = 16 cells. Unpaired t test (\*\*\*\**P*<0.0001).

As NuMA protein sequences are highly variable, an unbiased screen for LGN-interacting partners was used to identify and clone *Phallusia* NuMA (see Methods). Endogenous NuMA protein, recognized by an antibody made against a 190 amino acid portion of *Phallusia* NuMA (Fig. S1 D), is strongly enriched like LGN at the shared midline in B5.2 and B6.3 cell pairs (Fig. 3 G) and in live cells, NuMA-Venus accumulates at B5.2 cell contacts during mitosis (Fig. 3 H). Analysis of time lapse movies showed that the midline signal reaches a maximum at anaphase-telophase, when the signal intensity is twice as strong as that on the cortex near the other spindle pole (Fig. 3 H). In addition, in all cells NuMA protein is found on centrosomes during mitosis and increases at the cortex in interphase (Fig. S1 E), like the cyclic distribution described in mammalian cells (Seldin et al., 2013; Kotak et al., 2013, 2014; Zheng et al., 2014). Starting from the gastrula stage (5 hours post-fertilization), NuMA protein accumulates in interphase nuclei (Fig. S1, F and G), and both LGN and NuMA are found encircling the perimeters of apical cell surfaces (Fig. S1, G and H) similar to the vertebrate lateral belt (Peyre et al., 2011; di Pietro et al., 2016). In mitotic cells of the neuroepithelium (7 hours post-fertilization), LGN and NuMA are enriched on both lateral faces adjacent to spindle poles (Fig. S1 I) like in planar divisions of poliferating epithelia (Morin et al., 2007; Zheng et al., 2010; Tuncay and Ebnet, 2016; Nakajima, 2018).

These results show that in B5.2 and B6.3 cell pairs, LGN and NuMA localize at the shared midline cortex toward which spindles rotate. This localization is different from asymmetric cell divisions in sea urchin micromeres, *Drosophila* neuroblasts, or developing stratified epithelia, where LGN and NuMA accumulate on the exterior or apical surface promoting cell division perpendicular to the plane of the tissue (Poon et al., 2019; di Pietro et al., 2016; Williams and Fuchs, 2013; Tarannum et al., 2022). Rather, the topology of LGN and NuMA enrichment near the ascidian CAB structure suggests similarity to apical junctions (Fig. S1 J) which promote planar divisions in epithelia via lateral enrichment of LGN or NuMA (Bosveld et al., 2016; Tuncay and Ebnet, 2016; Higashi and Miller, 2017; Nakajima, 2018; Lechler and Mapelli, 2021).

### Inhibition of LGN/NuMA complex disrupts spindle position, unequal cleavage and bilateral symmetry

We next sought to inhibit the LGN/NuMA complex in order to assess its functions in spindle positioning and cleavage pattern. LGN is a maternal protein (Fig. S1 A) and its abundance was unaffected by the introduction of morpholino oligonucleotides. We thus used a dominant-negative approach to disrupt the LGN/NuMA cortical complex, by overexpressing the C terminal domain of LGN that attaches it to the plasma membrane or the portion of NuMA that binds to LGN (Fig. 4 A), both of which have been shown to have an inhibitory effect in other systems (Yu et al., 2002; Voronina and Wessel, 2006; Morin et al., 2007; Zheng et al., 2010; Peyre et al., 2011). In control embryos, B5.2 cell division occurs in a lateral-midline direction, with the smaller daughters connected at the midline and the larger daughters flanking them, to give the mirror symmetric planar UCD (green bars in Fig. 4 B and C). In B5.2 cells expressing the C terminal domain of LGN (LGN-Cter) or the LGN binding domain of NuMA (NuMA-LBD), cell division orientation was perturbed. The normal division axis was observed in only a fourth of cases, with the majority of B5.2 divisions (59% for LGN-Cter, 62% for NuMA-LBD) oriented parallel to the animal-vegetal axis (Fig. 4 B and D, blue bars). The remaining divisions (14%) took place along the anterior-posterior axis (Fig. 4 B and D, orange bars), which is never observed in control embryos.

**Figure 4.**
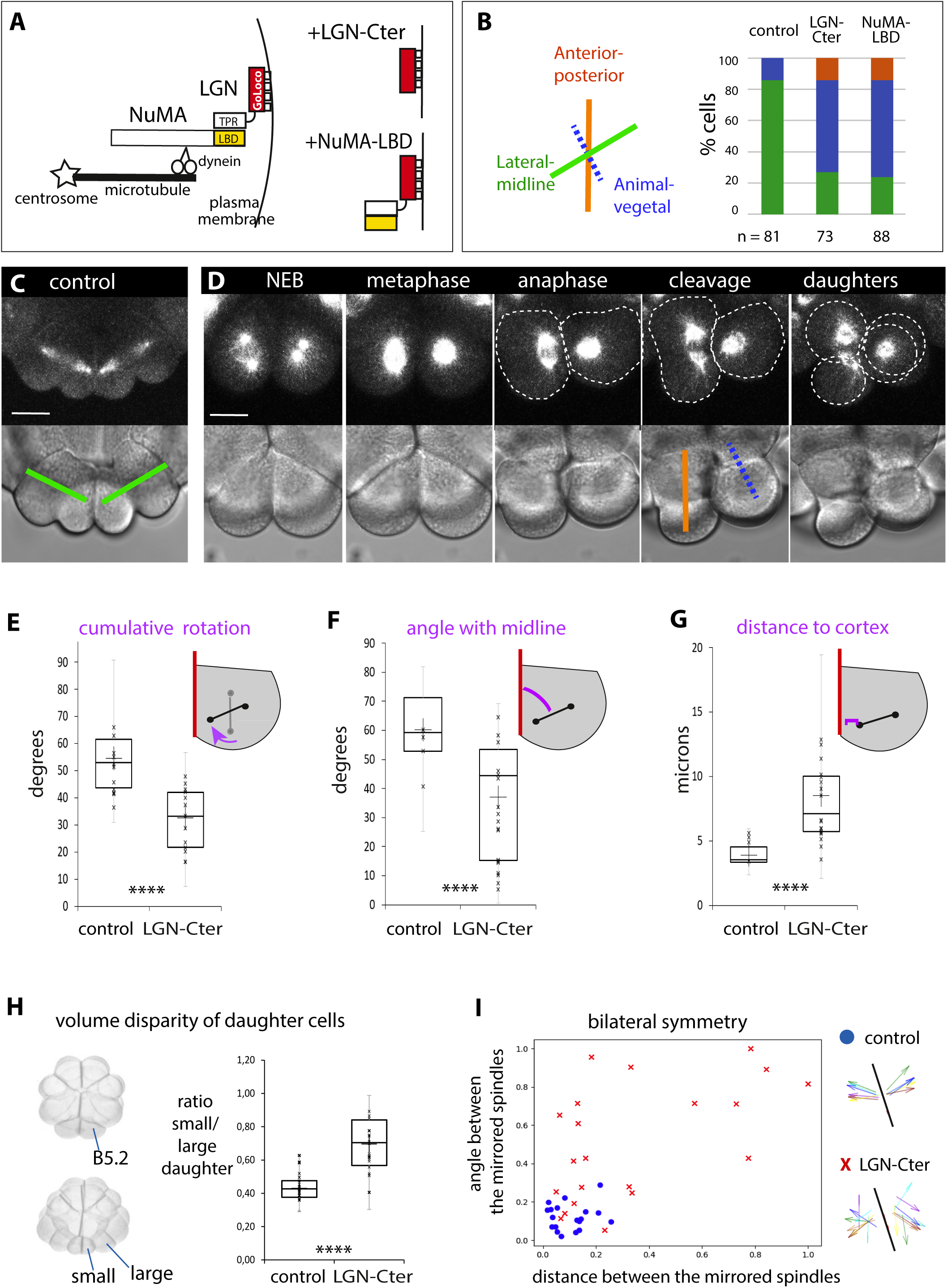
Inhibition of LGN/NuMA cortical complex disrupts spindle position, unequal cell division, and bilateral symmetry of ascidian germline B5.2 cells. A) Diagrams depicting the LGN/NuMA/dynein cortical complex (left) and two dominant-negative constructs used to disrupt it (right): LGN-Cter (in red) indicates the C terminal Goloco domains of LGN, and NuMA-LBD (in yellow) the LGN binding portion of NuMA. B) Orientation of B5.2 cell divisions in control embryos and embryos expressing dominant negative constructs. The colored axes (left) indicate 3 orthogonal directions: lateral-midline (green), animal-vegetal (blue), or anterior-posterior (orange). The bar graph (right) displays the percentage of each type of orientation and the number of B5.2 cell divisions analyzed (n) for three conditions. C) B5.2 cell divisions in a control embryo expressing EB3-Venus (in white) and H2B-RFP. Top shows spindle alignment at cleavage onset; bottom shows bright field image and the normal relative positions of daughter cells (green line = lateral-midline orientation). Scale bar = 20 microns. D) B5.2 cell divisions in an embryo expressing EB3-Venus (in white) and LGN-Cter dominant negative construct. Top shows spindle alignment throughout mitosis; bottom shows bright field images. One B5.2 cell divides in the anterior-posterior direction (dotted shapes on left cell and orange line); the other divides in the animal-vegetal orientation and generates 2 superimposed daughters (dotted shapes on right cell and blue line). Scale bar = 20 microns. E) Total spindle rotation (purple arrow in drawing), calculated as the angle in 3D between the spindle vector at NEB timepoint (gray line) and the spindle vector at anaphase timepoint (black line) in B5.2 cells expressing EB3-Venus and either H2B-RFP (control) or LGN-Cter construct. *N* = 36 cells for control, 30 for LGN-Cter. Unpaired t test (\*\*\*\**P*<0.0001). F) Angle in 3D of spindle with midline plane at anaphase (purple curve in drawing) in B5.2 cells expressing EB3-Venus and either H2B-RFP (control) or LGN-Cter construct. *n* = 36 cells for control, 44 for LGN-Cter. Mann-Whitney test (\*\*\*\**P*<0.0001). G) Distance between midline cortex and closest centrosome at anaphase (purple line in drawing) in B5.2 cells expressing EB3-Venus and either H2B-RFP (control) or LGN-Cter construct. *n* = 36 cells for control, 44 for LGN-Cter. Mann-Whitney test (\*\*\*\**P*<0.0001). H) Ratio of volumes of the daughter cells produced by B5.2 cell divisions in control embryos or embryos expressing dominant-negative construct LGN-Cter. The images on the left show a 16-cell stage embryo (top) and a 32-cell stage embryo (bottom). One B5.2 cell and its two daughter cells are indicated. *n* = 69 cells for control, 71 for LGN-Cter. Unpaired t test (\*\*\*\**P*<0.0001). I) Quantification of mirror symmetry. The graph shows the distance (x axis) and angle (y axis) between the two B5.2 spindles when projected mathematically onto the same side of the midline, normalized to maximum values. Each point represents one embryo expressing EB3-Venus and either H2B-RFP (control, blue circles) or LGN-Cter (red crosses). n= 18 embryos for control and 21 embryos for LGN-Cter. Unpaired t test for angle (\*\*\*\**P*<0.0001) and distance (\*\**P*=0.0024). Diagrams on right depict spindle orientations for pairs of B5.2 cells in representative embryos.The central black line corresponds to the midline plane and the same color is given to the two B5.2 spindles from one embryo.

Spindle movements were quantified in control embryos (Movie 3) and dominant-negative expressing embryos (Movie 4) by 3D tracking of the centrosome marker EB3-Venus. In B5.2 cells expressing LGN-Cter, the amount of spindle rotation (Fig. 4 E) and the final angle of the spindle with the midline at anaphase (Fig. 4 F) were significantly reduced compared to controls expressing H2B-RFP, and the centrosome-cortex distance at anaphase was doubled (average 8.76 microns compared to 4.17 microns in controls, Fig. 4 G). Consequently the volume disparity of the daughter cells produced by B5.2 cell division was decreased (ratio=0.7 ± .16 in LGN C-ter expressing embryos compared to 0.43 ± .08 in controls, Fig. 4 H). Finally, mirror-image spindle behavior was quantified by projecting the left and right spindles onto the same side of the midline plane and evaluating their overlap. In control embryos, the two B5.2 spindles show a small difference of position with values near 0 (blue circles in Fig. 4 I) while in LGN-Cter expressing embryos the spindles of two neighboring B5.2 cells superimpose only rarely (red crosses in Fig. 4 I) demonstrating a substantial loss of mirror symmetry. Injection of RNA encoding the LGN-Cter sequence from the closely related ascidian *Ciona intestinalis* led to misoriented cell divisions and spindle positioning defects very similar to those produced by the *Phallusia* LGN-Cter construct (Fig. S2 A-D). Overexpression of the N-terminal TPR domains of LGN however resulted in multipolar spindles and cytokinesis failure (Fig. S2 E-F) and likely disrupts the essential function of NuMA in spindle pole focusing (Merdes et al., 2000; Silk et al., 2009; Hueschen et al., 2017; Kiyomitsu and Boerner, 2021). Inhibition of dynein itself by overexpressing the dynactin component CC1 (Quintyne et al., 1999; Wuhr et al., 2010) led to a similar range of severe phenotypes (Fig. S2 G).

Overall these functional studies combined with invagination (Fig. 2) and localization (Fig. 3) results indicate that LGN/NuMA-mediated cortical pulling is responsible for spindle rotation, off-centering and mirror symmetry in the B5.2 germ line cells. This mechanism affords the great precision necessary for the geometric challenge of these particular cells, which must properly segregate the CAB and germ line determinants it contains into the smaller daughter cells while maintaining overall planar orientation of the embryonic spindles. In other cells of the ascidian embryo, spindle position is assured by less stringent mechanisms based on cell shape, cytoplasmic domains, or aster asymmetry (Negishi and Yasuo, 2015; Pierre et al., 2016; Costache et al., 2017; Dumollard et al., 2017; Godard et al., 2021). For instance, in the A6.2 cell pair, left-right mirror symmetry is only partial and spindle rotation occurs randomly clockwise or counterclockwise (Negishi and Yasuo, 2015). In contrast B5.2 cell pairs display near-perfect mirror symmetry, in both rotational direction and final spindle pole position. Interestingly, a transient polarized accumulation of LGN/NuMA is also implicated in spindle alignment in the bilaterally symmetric mitotic domains of the *Drosophila* gastrula (Camuglia et al., 2022).

### Cell cycle regulation of cortical pulling correlates with microtubules and not LGN/NuMA

Spindle positioning forces are regulated temporally as well as spatially. Recently we found that the strength of cortical pulling force is inversely related to CDK activity in 1- and 2-cell stage *Phallusia* embryos, i.e. cortical pulling is strong in interphase but diminishes upon entry to mitosis when spindles center (Rosfelter et al., 2024). We thus examined the temporal regulation of cortical pulling in 16-cell stage embryos using the invagination assay. In cytochalasin-treated embryos expressing fluorescent markers for plasma membrane and DNA, invaginations occur near spindle poles in all cells (Fig. 5 A). Time lapse microscopy showed that these invaginations do not appear at the time of NEB or metaphase but emerge around anaphase onset as indicated by chromosome separation (Fig. 5 A and B, also Fig. 2 D). The average time span from NEB to anaphase was 5.04±0.74 minutes followed closely by the average time from NEB to invagination at 5.5±-1.25 minutes (*n* = 32cells). To test this correlation, we delayed the metaphase-anaphase transition using the APC/C inhibitor MG132 and evaluated the timing of NEB, chromosome separation (anaphase), invagination appearance, and nuclear envelope reformation (NER) in timelapse movies. When anaphase onset and NER were delayed, invagination onset was also delayed (Fig. 5 C, Fig. S3 A and B). These results indicate that cortical pulling is not active during early mitosis (from prophase to metaphase) in cells of the 16-cell stage embryo and that it initiates at anaphase.

**Figure 5.**
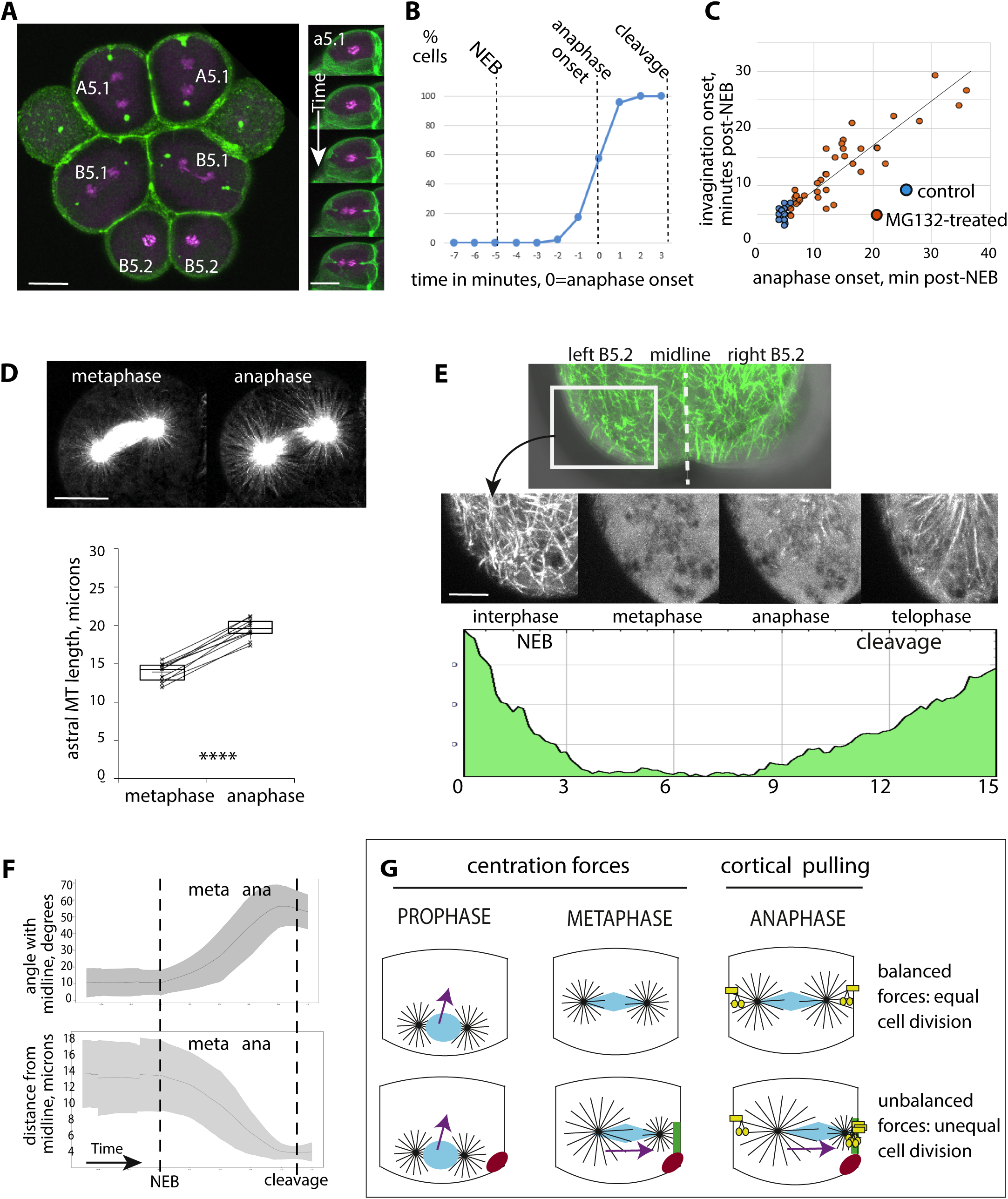
Cortical pulling initiates at anaphase upon microtubule lengthening. A) Cells from 16-cell stage embryos expressing PH-GFP (green) and H2B-RFP (magenta) and treated with cytochalasin. Labels in white indicate the names of the cells. Left: 4 vegetal cells in anaphase; green spots are the intracellular termini of membrane invaginations. The two B5.2 cells are in prometaphase. Right: images from timelapse video of a cell from the animal side. The time interval between images is 1 minute. Scale bars = 20 microns B) Percentage of cells in which invagination onset has occurred as a function of time; time scale is adjusted such that anaphase onset is at zero. Data from all cells of 16-cell stage embryos are grouped. *n* = 47 cells. C) Duration of time in minutes from NEB to anaphase (x axis) and from NEB to first invagination (y axis) in cells treated with cytochalasin (blue dots, *n*=32 cells) or cytochalasin and the APC/C inhibitor MG132 which delays cell cycle progression to anaphase (orange dots, *n* = 40 cells). Pearson correlation coefficient = 0.93. D) One cell from a 16-cell stage embryo expressing the microtubule binding protein Map7-GFP (in white) in metaphase and the same cell 3 minutes later in anaphase. The graph depicts the average length of 10 astral microtubules for 10 different cells. Lines connect the metaphase and anaphase values for the same aster. Paired t test (\*\*\*\**P*<0.0001). Scale bar = 20 microns. E) Top: pair of B5.2 cells expressing Ensconsin-GFP (green) to label microtubules. The dotted line indicates the midline. Cells are flattened against the coverslip and a plane closest to the plasma membrane is shown. 2^nd^ row: images from timelapse video of the boxed region, showing cortical microtubules (white) at the indicated cell cycle stages. The graph depicts the intensity of fluorescent signal of the selected region (y axis) over time in minutes (x axis). a.u.=arbitrary units. NEB=nuclear envelope breakdown. F) Angle of spindle with midline plane (top) and distance of spindle center from midline plane (bottom) as functions of time (x axes). The data from 26 films are interpolated to a normalized time scale with 0 = NEB and 1 = cytokinesis. The central lines are the mean, and the gray curves are the standard deviation. G) Model depicting how forces governing spindle position are influenced by the cell cycle in the ascidian embryo. The top row represents an equally dividing cell; the bottom row indicates one B5.2 cell containing the CAB (in red) and enrichment of LGN and NuMA on the midline (in green). Astral microtubules are in black, and the nucleus or spindle in blue. Arrows indicate direction of movement of the spindle center. Upon entry to mitosis (prophase), the two asters and nucleus move to the center in all cells, then after nuclear envelope breakdown the spindle either remains centered if forces are balanced (top row), or shifts off-center due to unbalanced forces (bottom row). The yellow shape (anaphase column) represents dynein tethered to the membrane exerting pulling force, either equally on both sides (top row) or stronger at the midline side (bottom row).

This switch-like regulation of cortical pulling, from off in metaphase to on in anaphase, could be due to the cyclic localization of NuMA which moves from spindle poles to the cell cortex upon dephosphorylation in anaphase (Gehmlich et al., 2004; Selden et al., 2013; Kotak et al., 2013; Kotak et al., 2014; Zheng et al., 2014; Gloerich et al., 2017; Lee et al., 2018; Kiyomitsu and Boerner, 2021; Fig. S1 E). However our finding in B5.2 cells that cortical pulling is still delayed until anaphase even if LGN/NuMA are already cortically enriched in metaphase (Fig. 3) indicates that some other component required for force generation is missing in metaphase. We hypothesize that reduced cortical pulling during early mitosis is due to changes in MTs, which become shorter and more dynamic when CDK activity is high (Belmont et al., 1990; Verde et al., 1990; Verde et al., 1992; Cassimeris, 1999; Rusan et al., 2001). We examined live 16 cell stage *Phallusia* embryos and found that in all cells MTs fill the cytoplasm during interphase then concentrate around the spindle during mitosis (Fig. S3 C, Movie 5) as previously observed (Prodon et al., 2010; Costache et al., 2017; Dumollard et al., 2017). Measurement of astral MT length showed an increase of 29% between metaphase and anaphase (from 13.87 ± 1.24 microns to 19.54 ± 1.33 microns, Fig. 5 D). The minimum distance from centrosome to cell surface in these same cells was measured at 17.50 ± 3.19 microns, indicating that indeed on average metaphase MTs are too short to reach the cortex. Analysis of z slices just near the cell cortex revealed that MT density at the cell surface is high in interphase, decreases dramatically upon entry to mitosis, then increases at anaphase prior to cytokinesis (Fig. 5 E, Fig. S3 D). These results indicate that the downregulation of cortical pulling at the onset of mitosis is due to the shortening of microtubules which would prevent their interaction with cortical dynein, and conversely that the lengthening of MTs at anaphase onset allows cortical pulling to be effective.

In the germ line pair of cells, a substantial amount of spindle rotation and migration occurs in metaphase prior to the onset of cortical pulling at anaphase (Fig. 1 B, Fig. 5 F, Prodon et al., 2010; Costache et al., 2017; Dumollard et al., 2017). We previously showed that the two mitotic asters in the ascidian germ line cells are different in size, due to enrichment of the MT depolymerase Kif2/MCAK in the CAB (Costache et al., 2017). We therefore propose that the metaphase movement in B5.2 cells could be caused by cytoplasmic pulling applied to non-identical asters, as opposed to its classical role of centering symmetrical spindles. Then during anaphase, cortical pulling enhances spindle pole migration due to a local accumulation of LGN/NuMA at the midline contact site.

Overall our results point towards a 2 step mechanism whereby the spindle is moved by different directional forces during distinct phases of the cell cycle: starting at prophase the reduction of cortical pulling and of microtubule length frees asters from their cortical tethers allowing centration forces to dominate during metaphase (via cytoplasmic pulling or pushing from astral growth), then in anaphase astral MTs are captured by cortical pullers thus anchoring spindle poles and stabilizing their positions (Fig. 5 G). This temporal separation of cytoplasmic and cortical pulling may explain how opposing forces can cooperate for spindle positioning. A similar 2 step regulation is expected in large cells of early embryos where astral MTs do not reach the cortex in metaphase but lengthen considerably at anaphase (Strickland et al., 2005; Foe and VonDassow, 2008; Wuhr et al., 2010; Mitchison et al., 2012; Rieckhoff et al., 2020; Xie et al., 2022; Xie et al., 2025). Even in smaller cells of yeast (Zucca et al., 2023) or human cell lines (Singh et al., 2021), destabilization of astral MTs in early mitosis by CDK activity is necessary for correct spindle positioning. Other mechanisms have been described to reduce or override metaphase cortical pulling for equal cell division (Kiyomitsu and Cheeseman, 2012; Middelkoop et al., 2024) or to harness it for unequal cell division (Labbé et al., 2004; Wu et al., 2024). Future work taking into account the temporal as well as spatial regulations of motor complexes and microtubule dynamics will uncover the diverse combinations which achieve accurate spindle position in each cell type.

## Materials and methods

### Ascidian egg fertilization and embryo culture

All experiments were performed with the ascidian *Phallusia mammillata*, a marine chordate. Adult *Phallusia* were collected at Sète (Etang de Tau, Mediterranean coast, France) or Roscoff (Brittany, France) and maintained for several months in aquaria at the Centre de Ressources Biologiques (CRB) of the Institut de la Mer à Villefranche (IMEV). Following dissection of the adult hermaphrodite, gametes were collected with pipets from the oviduct and spermiduct. Eggs were de-chorionated by treatment with 0.1% trypsin (Sigma-Aldrich, T9201) in micro-filtered seawater (MFSW) buffered with 5 mM TAPS pH 8.2 ([tris (hydroxymethyl) methylamino] propanesulfonic acid) for 1-2 hours. Because dechorionated eggs adhere to glass and plastic, all dishes and pipettes are coated with agarose or “GF” (a mixture of 0.1% gelatin and 0.1% formaldehyde) (Sardet et al., 2011; McDougall et al., 2015). To fertilize, sperm was activated by dilution into pH 9.5 sea water for 10 minutes then added to eggs at a ratio of ∼1: 100. Once eggs changed shape which indicates fertilization, they were transferred to fresh MFSW to prevent polyspermy. Embryos were then cultured in GF-coated dishes at 18°C until the desired stage.

### Microinjection

Injection needles were made from thin wall glass capillaries without filament, inner diameter 0.78 mm outer diameter 1 mm (GC100T-10, Harvard Apparatus) using a Narishige PN-30 horizontal puller. Needles are placed in a HI-7 needle holder which receives pressure from a Narishige IM300 injection box attached by oil-filled tubing to a Mecafer air compressor. The needle is controlled by a three-axis hydraulic micromanipulator (Narishige MMO-203) mounted on a Nikon DiaPhot inverted microscope fitted with 10X and 20X objectives. Dechorionated eggs were placed in horizontal glass injection chambers filled with MFSW (Sardet et al., 2011; Chenevert et al., 2024). Needles were front-filled from injection tubes prepared from glass capillaries (GC100T-10, Harvard Apparatus) containing 1 µl RNA flanked by 1 µl layers of mineral oil (Sigma). The concentration of RNA in the needle was 1-5 μg/μl and the amount injected was 2-5% egg volume as estimated by the small clearing in the cytoplasm. Injected eggs were left at 18°C for several hours or overnight before fertilization and imaging or collection for western blot.

### Cloning of ascidian LGN and NuMA

Homology searches identified the sequences corresponding to LGN from *Phallusia mammillata* (GenBank CAB3250827.1) and *Ciona intestinalis* (NCBI XP_018672345.1). Full length clones were selected from arrayed cDNA libraries made from *Phallusia* RNA (GenBank accession number PX794811) or *Ciona* RNA (GenBank accession number PX794812). NuMA protein sequences are highly variable thus difficult to identify by homology, but searches for proteins with similar domain organization identified one potential candidate (GenBank accession number CAB3265828.1). This same gene was cloned as the strongest interactor in a yeast two hybrid screen for partners of *Phallusia* LGN (PX794810), confirming its identity as *Phallusia* NuMA.

For the yeast two hybrid screen, a library was constructed with cDNA of *Phallusia* embryos fused to the GAL-4 activation domain in the vector pEXP-AD502, using the ProQuest system (Invitrogen). This *Phallusia* library was transformed into yeast strain MAV203 and screened with a plasmid containing *Phallusia* LGN coding sequence fused to the GAL-4 DNA binding domain in the vector pDBLeu. Yeast colonies positive for the His3 reporter indicating GAL4-dependent transcription were selected according to ProQuest protocols (Invitrogen). Secondary screening for two other reporter genes, Ura3 and lacZ, resulted in three strongly positive clones, all of which contained the C terminal region of the open reading frame corresponding to GenBank accession number PX794810, beginning at amino acid 816 or 900. The results of this methodology both confirmed the identity of ascidian NuMA coding sequence (GenBank accession number PX794810) and specified the LGN binding domain of *Phallusia* NuMA (denoted NuMA-LBD in the text).

### Constructs and RNA synthesis for microinjection

To label cellular organelles, fluorescent reporter sequences (encoding GFP, Venus, RFP, dTomato) were added as in-frame fusions to the C termini of marker proteins, as described in previous publications (McDougall et al., 2015) : Histone H2B (mouse sequence, GenBank accession number BI105582.1) for chromosomes (Prodon et al., 2010; Dumollard et al., 2017), EB3 (human sequence, GenBank accession number AY893969.1, gift from P. Lenart) for centrosomes (Rosfelter et al., 2024), Map7 (N terminal fragment of mouse sequence, GenBank accession number BC052637) and Ensconsin (human sequence, GenBank accession number NP_003971.1) for microtubules (Prodon et al., 2010; Costache et al., 2017), and PH domain (PIP_2_ binding domain from human PLCδ1) for plasma membrane (Prodon et al., 2010). The NLS sequence to visualize nuclei is ATGACTGCTCCAAAGAAGAAGCGTAAGGTA which encodes the nuclear localization signal from SV40 large T antigen (MTAPKKKRKV). Venus-LGN is Venus fluorescent protein fused to the N terminus of full length LGN from ascidian *Ciona intestinalis* (M1-E644, GenBank accession number PX794812). NuMA-Venus is Venus fluorescent protein fused to the C terminus of full length NuMA from *Phallusia mammillata* (M1-K1517, GenBank accession number PX794810). For dominant negative mutant constructs, portions of LGN or NuMA coding sequences were prepared as “no-tag” constructs, lacking fluorescent reporter additions. These were *Phallusia* LGN-Cter = G451-E653, comprising the 4 “Goloco” domains; *Ciona* LGN-Cter = E453-E644, comprising the 4 “Goloco” domains; *Ciona* LGN-Nter = M1-G451, comprising the 6 “TPR” domains; *Phallusia* NuMA-LBD = D900-L1205, comprising the LGN binding domain. The dynein inhibitor CC1 is a 334 amino acid portion of the coding sequence of p150 glued protein, *Ciona intestinalis* dynactin subunit 1 (S193-S526, NCBI reference sequence XP_018673000.1).

All coding sequences of interest were cloned into vectors derived from pRN3 which harbors a promotor for T3 RNA polymerase and oligo dT sequences for the insertion of a polyA tail. Following linearization of the plasmids with Acc65i or Sfi1, RNAs were transcribed using T3 RNA polymerase and 5’ capped with the mMessage mMachine kit (Ambion).

### Generation of specific antibodies to LGN and NuMA

The full-length open reading frame of *Phallusia* LGN (M1-E653) was fused to a 6-histidine tag at the C terminus and cloned into the vector pET11. Expression in bacteria was induced with IPTG and the LGN fusion protein was purified on a column of nickel resin (Protino Ni-NTA agarose) and eluted with imidazole according to methods of the manufacturer (Macherey-Nagel). Immunization of mice was performed by Covalab (France) and sera obtained after 67 days were used directly without further purification. Antibodies to *Phallusia* NuMA protein were produced in rabbits by Proteogenix (France) using as antigen the polypeptide corresponding to amino acids I67-N257 of *Phallusia* NuMA (GenBank accession number PX794810). Sera obtained after 51 days were purified against the Numa antigen by Proteogenix before use.

### Immunoblotting

Western blots were used to evaluate specificity of our antibodies and to assess levels of endogenous LGN and NuMA. Live eggs or embryos were collected in a small volume of MFSW and mixed with the same volume of 2× Laemmli sample buffer [100 mM Tris-HCl (pH 6.8), 4% SDS, 0.2% Bromophenol Blue, 20% glycerol and 200 mM dithiothreitol). Samples were heated for 5 min at 95°C and then stored at −20°C. Proteins were separated by electrophoresis on 8% or 10% SDS-polyacrylamide gels and transferred to a nitrocellulose membrane (Amersham). Membranes were stained briefly with Ponceau-S, then washed into TBS-Tw (20 mM Tris Base and 150 mM NaCl plus 0.1% Tween-20), and blocked for at least 1 hour in TBS-Tw containing 5% dry milk powder. Immunoblots were then incubated in primary antibodies diluted 1:1000 overnight at 4°C. After antibody incubation, membranes were washed three times with TBS-Tw and incubated with an appropriate horseradish peroxidase-conjugated secondary antibody (Jackson ImmunoResearch) at 1:10,000 dilution in TBS-Tw + 5% milk powder for 2 hours at room temperature. After three washes with TBS-Tw, signal detection was carried out using the SuperSignal West Pico chemiluminescent substrate (ThermoFisher Scientific) and images were acquired with a Fusion FX system (Vilber Lourmat, Collegien, France).

### Immunofluorescence

For all antibody labelling, embryos were fixed in 100% methanol which had been pre-chilled at −20°C and then stored at -20°C. Samples were rehydrated by three washes in PBS containing 0.1% Tween (PBS-Tw), then blocked in PBS-Tw containing 1% BSA (bovine serum albumin, Merck) for 1 hour at room temperature. Primary antibodies were added in the blocking solution and incubated overnight or longer at 4°C. Primaries and dilutions used were anti-tubulin DM1a (mouse Sigma-Aldrich) at 1:500, anti-gamma tubulin GTU88 (mouse Sigma-Aldrich) at 1:200, anti-aPKC (rabbit Santa Cruz Biotechnology sc-216) at 1:200, anti-Par3 (rabbit, made against *Drosophila* Bazooka gift from Andreas Wodarz) at 1:300, anti-LGN (mouse, this study) at 1:200, anti-NuMA (rabbit, this study) at 1:200. After incubation, samples were washed three times in PBS-Tw and appropriate fluorescent secondary antibodies (Jackson ImmunoResearch) were added at a dilution of 1:200. After further incubation at room temperature for 1-2 hours, samples were washed in PBS-Tw, labelled with 5 µg/ml Hoechst 33342 (Sigma-Aldrich) for 10 minutes, washed twice more in PBS-Tw and finally resuspended in anti-fading agent CitiFLuor AF1 (Electron Microscopy Services).

For 3D reconstruction of the LGN midline domain (Movie 2), a confocal Z stack of a pair of B5.2 cells labelled with anti-LGN, anti-Par3, and Hoechst was segmented with Fiji software (Schindelin et al., 2012) and adjusted with the threshold function. A shape mesh was extracted in Python using Marching Cube algorithm and Laplacian smoothing and visualized with Napari software (Chiu et al., 2022).

### Image acquisition

Imaging was performed with a Leica TCS SP8 inverted confocal microscope fitted with a 40×/1.1NA water immersion objective and 408, 488 and 552 nm lasers and Leica application suite acquisition software (LASX), or with a Zeiss Axiovert 200M inverted microscope fitted with a 40×/0.75NA objective and equipped with a CoolSnap Kino camera (Photometrics) or ORCA digital Fusion camera (Hamamatsu) and Metamorph acquisition software (Molecular Devices).

For mounting of fixed immuno-stained samples, embryos were placed in a drop of antifadent solution CitiFluor AF1 (Electron Microscopy Services) on glass slides. To prevent embryo compression or breakage, coverslips were elevated by application of small spacers of modelling clay at the corners as described (Sardet et al., 2011) and sealed with nail polish to prevent evaporation.

For mounting of live samples, embryos were placed in a drop of MFSW in GF-coated glass bottom dishes (Cellvis) secured with a coverslip as described (Chenevert et al., 2024) or on a GF coated slide and covered with a GF coated coverslip which was elevated and sealed with 4 lines of silicone vacuum grease (Dow-Corning) as described (Sardet et al., 2011). Full Z scans of the relevant cells were acquired in both fluorescent and brightfield channels with Z step 1-2 microns and time interval 30-120 seconds, depending on the number of stage positions to be filmed. All live imaging experiments were performed at 18–19 °C.

### Measurement of degree of spindle shift in fixed samples

Embryos were fixed in cold (-20°C) methanol every 10 minutes from 2 to 4 hours post-fertilization and labelled for centrosomes (anti-gamma tubulin GTU88), a CAB marker (aPKC or Par3), and DNA (Hoechst). Confocal stacks of B4.1, B5.2, or B6.3 cells in anaphase were acquired with a Z-step of 1 micron. To view the cell plane containing the spindle, each Z stack was rotated using the Oblique Slicer tool of Imaris software (BitPlane) until both centrosomes were in the same plane of view. The distances from the center of the spindle to the cell periphery along the spindle axis were measured to give (a) the shortest distance and (b) the longest distance. The ratio a/b defines the degree of spindle shift.

### Invagination assay to visualize cortical pulling

Plasma membrane was labelled by microinjection of synthetic RNA encoding PH-dTomato or PH-GFP or by addition of the lipophilic dye Cell Mask Green (Thermo Fisher Scientific C37608) or Cell Mask Orange (Thermo Fisher Scientific C10045) at 1/1000 dilution in MFSW. Cytochalasin B (Sigma C6762) was kept frozen as a 10 mM stock solution in DMSO and added to embryos at 5-10 µg/ml just prior to imaging. Image stacks were acquired at a z step of 1 micron and time interval of 1-2 minutes. As invaginations can occur anywhere on the cell surface not just in the plane of view, the timelapse videos were analyzed with Imaris software (Bitplane) in surpass mode 3D view which allows images to be visualized in all orientations and observation of a maximum number of invaginations. The proteasome inhibitor MG132 (Sigma 474790) was kept frozen as a 20 mM stock solution in DMSO and added to embryos at 10 µg/ml at the 4-cell stage, followed by cytochalasin treatment at the 16-cell stage and imaging.

### Quantification of cortical LGN and NuMA

Fluorescence intensity of LGN or NuMA was measured on maximum intensity projections of confocal Z-slices comprising labelled chromosomes and the CAB using Fiji image analysis software (Schindelin et al., 2012). For fixed B5.2 and B6.3 cells labelled with anti-LGN (Fig. 3 E and F), 3 lines of length 10 microns and pixel width 3 were drawn perpendicular to the apical (outside) surface of the embryo: one along the midline starting at the CAB, one along the lateral membrane opposite to midline, and one into the center of the cell in order to evaluate the cytosolic background. For each of the 3 lines (midline, lateral, or background), the fluorescence intensity was obtained using plot profile function. All values corresponding to the first 6 microns were averaged and used to calculate the ratios midline/background and lateral/background. To evaluate intensity along the midline as a function of distance from apical (Fig. S1 B), fluorescent intensity values were separated into 1 μm increments starting from the CAB side.

For films of live B5.2 cells expressing NuMA-Venus (Fig. 3 H), at each timepoint 2 lines of 7-pixel thickness were drawn perpendicular to the midline cell contact or the opposite cell contact. The peak intensity values obtained with the Fiji plot profile function were designated midline or lateral (for opposite cell contact) and divided by cytosolic background (average of 8 values). To combine data from different timelapse movies, a normalized time scale was developed as follows. Analysis of control B5.2 cells expressing a fluorescent DNA marker (Histone H2B) showed that the average duration from NEB to cleavage onset is 6.52 minutes (391 ± 55 seconds, n= 38) and chromosome separation begins 4.82 minutes after NEB (289 ± 52 seconds, n=24). Thus if the timing of NEB = 0 and cleavage = 1, then anaphase onset = 0.73. For each NuMA-Venus film, the timepoints between NEB and cytokinesis onset were grouped into sequential thirds, designated prometaphase, metaphase and anaphase, and fluorescent intensity values within each group were averaged.

### Extraction of spindle pole coordinates from live timelapse data

To quantify spindle movements, *Phallusia* eggs were injected with a mixture of two synthetic RNAs encoding centrosome marker EB3-Venus and DNA label H2B-RFP (as control) or with RNAs encoding EB3-Venus and LGN-Cter dominant negative construct. These micro-injected eggs were incubated overnight at 18°C then fertilized and mounted on GF-coated slides 90 minutes later at the 4-cell stage in order to obtain a standardized orientation with the animal-vegetal axis and the midline plane perpendicular to the XY plane of view (McDougall et al., 2015). The slides were maintained in humidified chambers at 18°C for a further hour until the 16-cell stage then subjected to multichannel video microscopy. For each embryo, Z stacks were set to cover the vegetal side which contains the germ line B5.2 cells, and the resultant movies were analyzed with Fiji software (Schindelin et al. 2012). XYZ coordinates for centrosomes and points on the cell surface were obtained by 3D manual tracking using the Fiji plugin MTrackJ (Meijering et al., 2012). The timepoints corresponding to mitotic events were determined visually from the H2B signal for anaphase onset or from brightfield stacks for nuclear envelope breakdown and cytokinesis onset.

### Calculations of spindle geometry

The angle change of a spindle over time (Fig. 4 E and Fig. S2 A) is the angle between the spindle at the timepoint of NEB (vector *S_n_*) and the same spindle at anaphase (vector *S_a_*):

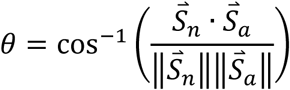

The angle between a spindle and the midline plane (Fig. 4 F and Fig. S2 B) is a function of the dot product of the spindle vector and the normal to the midline plane:

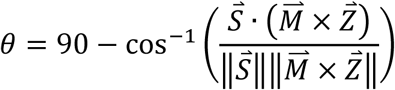

where S is the spindle vector directed from the internal centrosome (closest to the midline) to the external centrosome (lateral or farther from the midline), M is the midline vector directed from a point on the midline toward the apical spot where the two CABs meet, and Z is the z axis which is also part of the midline plane.

The distance d from a centrosome (coordinates *x*1, *y*1, *z*_1)_) to the closest point on the cortex (coordinates *x*_2_, *y*_2_, *z*_2_)(Fig. 4 G and Fig. S2 C) is:

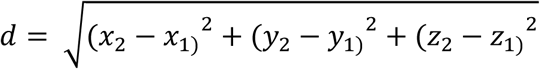

To quantify the degree of mirror symmetry (Fig. 4 I, Fig. S2 D), an XY rotation was performed to align the midline vector M with the y axis. As a result, the midline plane coincides with the YZ plane of imaging and the embryonic axes are aligned on the grid of coordinates, with x axis = midline-lateral, y axis = anterior-posterior, and z axis = animal-vegetal. The anaphase spindles in the two neighboring B5.2 cells, designated vector 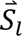 and vector 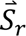 for Spindle left and Spindle right respectively, were placed on the same side of the midline plane YZ by replacing each x position value by its absolute value. The distance d between the two spindle centers C_l_ and C_r_ and the angle φ between vectors 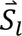 and 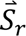 were then calculated:

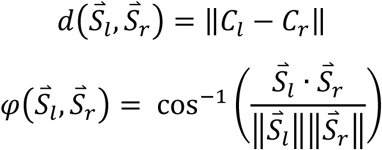

If the two spindles superimpose which is the case for perfect mirror symmetry, d and φ are zero. For graphical representation the distance and angle values were normalized to the maximum value. To represent the geometric configuration of spindles in two neighbor cells as colored arrows, normalized vectors in 3D space were generated in Python.

### Cell volume calculation

From time lapse videos the Z-stack corresponding to completion of B5.2 cytokinesis, when newborn cells appear roughly spherical, was selected. Measurement of surface area (A) was obtained for each of the two daughter cells, B6.3 and B6.4, by tracing their perimeters at the equatorial plane using Fiji segmented line tool. The cell radius (*r*) and volume (*v*) were estimated from the area (A):

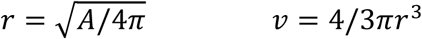

The ratio of volumes as a function of the two areas is:

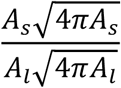

where *A*s and *A*l are the areas of the smaller and larger daughter cells, respectively.

### Quantification of astral microtubule length and abundance

Images of 16 cell stage embryos expressing MAP7-GFP or Ensconsin-GFP were acquired on a Leica SP8 confocal microscope, with a Z step of 1 micron. For each timepoint, a maximum projection was made of the 3 Z planes containing the centrosome and thus in-focus astral microtubules. The lengths of the 10 longest MTs visible were measured in a given aster in metaphase and in the same aster several minutes later in anaphase, using Fiji line measurement tool. The values for asters from 10 different cells are displayed on Fig. 5 D. To visualize surface microtubules (Fig. 5 E), confocal images of a 1 micron thick optical section just above the plasma membrane were acquired at a rate of 8 seconds/image. After thresholding of the xyt stack, the fluorescent intensity of microtubule signal on the surface was quantified with Fiji measure stack function (Schindelin et al. 2012).

### Statistical analysis

Statistical analysis was performed with Excel (Microsoft) and Prism 8 (GraphPad). Normality was evaluated with a Shapiro-Wilk test except for Fig. 3 E and Fig. S2 A where a Kolmogorov-Smirnov test was performed. The statistical tests applied and *P* values obtained are indicated in each figure legend. Box and whisker plots represent the distribution of the data using quartiles. Q1 and Q3 form the limits of the box and the median Q2 is indicated by a bold line.

The whiskers represent the range in which most values are found, whereas the values outside represent the outliers. The small line or cross in the box indicates the mean.

## Acknowledgements

We thank Stefania Castagnetti, Benoit Godard, Benjamin Lacroix, and Gerard Pruliere for valuable discussions, Andreas Wodarz for the gift of the Par3 antibody, and Stefania Castegnetti for critical reading of the manuscript. We are grateful to Faisal Bekkouche, Kelly Daniel, Philippe Dru, Margaux Failla, Laurent Gilletta, Celine Hebras, and Sebastien Schaub for technical assistance. The project was funded by the Agence Nationale de la Recherche grants ‘‘MorCell’’ (ANR-17-CE 13-0028) and “InvBlastula” (ANR-22-CE13-0036-01). We thank the Service Aquariologie of CRB (IMEV-FR3761) and the Imaging Facility (PIM, a member of the MICA microscopy platform) which are supported by the EMBRC-France infrastructure network (Grant ANR-10-INBS-02). The authors declare that they have no conflict of interest.

## Author contributions

J. Chenevert and A. Rosfelter performed, analyzed and interpreted the experiments. J. Chenevert, R. Dumollard and A. McDougall conceived and designed the project. S. Caballero-Mancebo and D. Gonzales-Suarez contributed to data analysis. L. Besnardeau constructed molecular tools. V. Costache, P. Stolz, R. Dumollard, and A. McDougall contributed experimental data. J. Chenevert, R. Dumollard, and A. McDougall wrote the manuscript with contributions from A. Rosfelter and S. Caballero-Mancebo.

## Figures legends

**Figure S1.**
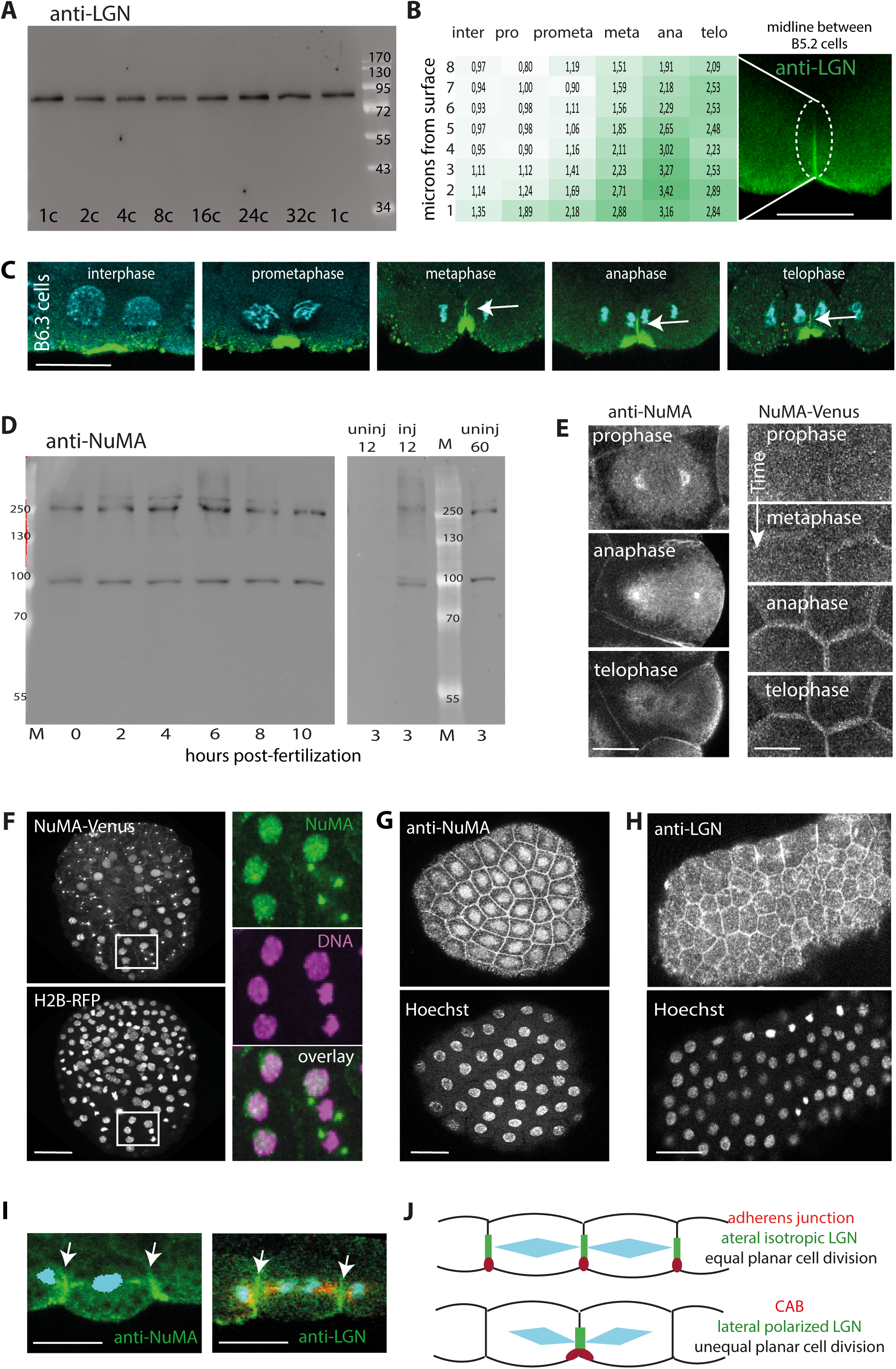
Localizations of LGN and NuMA in ascidian embryos. (Related to Figure 3) In all panels, scale bars = 20 microns. A) Immunoblot of *Phallusia* protein extracts probed with anti-LGN antibody. Each lane was loaded with protein from 40 embryos lysed at the indicated stages (1-, 2-, 4-, 8-, 16-, 24-, or 32-cell stages). The size of molecular weight markers in kilo-daltons is written on the right. The calculated molecular weight of *Phallusia* LGN is 73 kDa. B) Table indicating the strength of anti-LGN label as functions of distance from the apical surface and phase of mitotic progression. The ratio of fluorescence intensity along the midline over cytoplasmic background was evaluated in 1 micron increments between the apical face (bottom) and 8 microns inward toward the basal side (top). The values are color-coded: highest intensity is bright green and lowest is white. LGN accumulation along the midline both initiates and achieves its highest level near the CAB which is situated apically. C) Pairs of B6.3 cells fixed at the indicated phase of the cell cycle and stained with LGN antibody (green) and Hoechst (cyan). White arrows indicate LGN signal at the midline. D) Immunoblot of *Phallusia* protein extracts probed with anti-NuMA antibody. Left: each lane was loaded with equal amounts of embryos lysed at the indicated time of development (in hours post-fertilization). The calculated molecular weight of *Phallusia* NuMA is 175 kDa. Two major bands of 250 and 110 kDa are detected at all stages. Right: The same bands of 250 and 110 kDa are enriched in embryos overexpressing NuMA. Lanes were loaded with protein from 12 uninjected embryos (“uninj”), 12 embryos overproducing *Phallusia* NuMA coding sequence (“inj”=injected), or 60 uninjected embryos, all harvested at 3 hours post-fertilization. M: molecular weight marker lane; sizes are indicated on both sides in kilo-daltons. E) Changes in NuMA protein localization during mitosis in cells from 16-cell stage embryos. Left: confocal plane containing centrosomes from cells labelled with anti-NuMA antibody (white); right: surface view of cells expressing NuMA-Venus (white). Cell cycle stages are indicated. F) Gastrula stage embryo expressing NuMA-Venus (top, green in right panel) and H2B-RFP (bottom, pink in right panel). Right panels are enlargement of the boxed region showing 4 cells in interphase (NuMA in nuclei) and 2 cells in mitosis (NuMA on centrosomes). G) Gastrula stage embryo fixed and stained with anti-NuMA (top) and Hoechst for DNA (bottom). H) Neurula stage embryo fixed and stained with anti-LGN (top) and Hoechst for DNA (bottom). I) Mitotic cells from neurula stage embryos immuno-labelled with anti-NuMA (green, left) or with anti-LGN (green, right) and anti-tubulin (red) and Hoechst (blue). Arrows indicate enrichment of NuMA or LGN at lateral cell contacts adjacent to both spindle poles. J) Drawing depicting neighboring cells undergoing equal planar division (epithelia, top) or unequal planar division (ascidian germ line, bottom). Red ovals indicate the position of adherens junctions or CABs; green bars indicate localized cortical LGN and NuMA; blue diamond shapes represent spindles.

**Figure S2.**
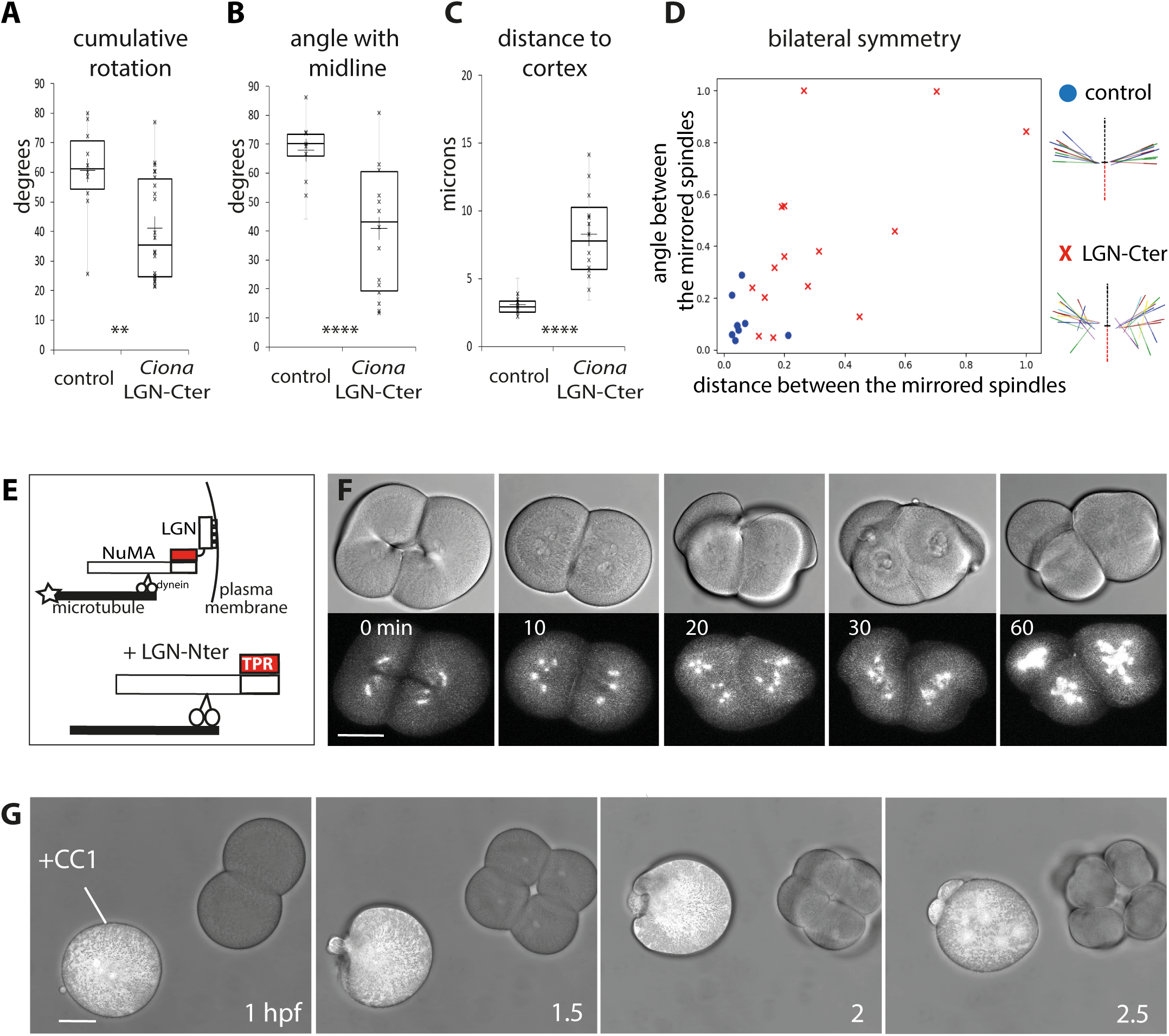
Phenotypes generated by disruption of LGN, NuMA, or dynein. (Related to Figure 4) A) Total spindle rotation calculated as the 3D angle between the spindle vector at NEB and the spindle vector 4-5 minutes later at anaphase in B5.2 cells expressing EB3-Venus and either H2B-RFP (control) or *Ciona* LGN-Cter construct. *n* = 10 cells for control, 22 for *Ciona* LGN-Cter. Unpaired t test (\*\**P*=0.0052). B) Angle of spindle with midline plane at anaphase in B5.2 cells expressing EB3-Venus and either H2B-RFP (control) or *Ciona* LGN-Cter construct. *n* = 16 cells for control, 30 for *Ciona* LGN-Cter. Mann-Whitney test (\*\*\*\**P*<0.0001). C) Distance between midline cortex and closest centrosome at anaphase in B5.2 cells expressing EB3-Venus and either H2B-RFP (control) or *Ciona* LGN-Cter construct. *n* = 16 cells for control, 30 for *Ciona* LGN-Cter. Unpaired t test (\*\*\*\**P*<0.0001). D) Quantification of mirror symmetry. The graph shows the distance (x axis) and angle (y axis) between the two B5.2 spindles when projected mathematically onto the same side of the midline, normalized to maximum values. Each point represents one embryo expressing EB3-Venus and either H2B-RFP (control, blue circles) or *Ciona* LGN-Cter (red crosses). *n* = 8 embryos for control and 15 for LGN-Cter. Unpaired t test for angle (\**P*=0.0122) and Mann-Whtney test for distance (\*\*\*\**P*<0.0001). The diagrams depict spindle orientations for pairs of B5.2 cells in representative embryos. The central dotted line corresponds to the midline and the same color is given to the two B5.2 spindles from one embryo. E) Diagrams depicting the LGN/NuMA/dynein cortical complex (top) and overexpression of the N-terminal TPR domains of LGN (in red) which bind to NuMA (bottom). F) Bright-field (top row) and fluorescent (bottom row) images from a timelapse video of an embryo expressing LGN-Nter and the microtubule marker EB3-Venus (in white). Time in minutes is indicated. Scale bar = 50 microns. G) Overlay of brightfield and fluorescent signals from two embryos fertilized and filmed at the same time, one uninjected control and the other expressing CC1-Venus (in white) which inhibits dynein. Time is indicated as hours post-fertilization (hpf). Scale bar = 50 microns.

**Figure S3.**
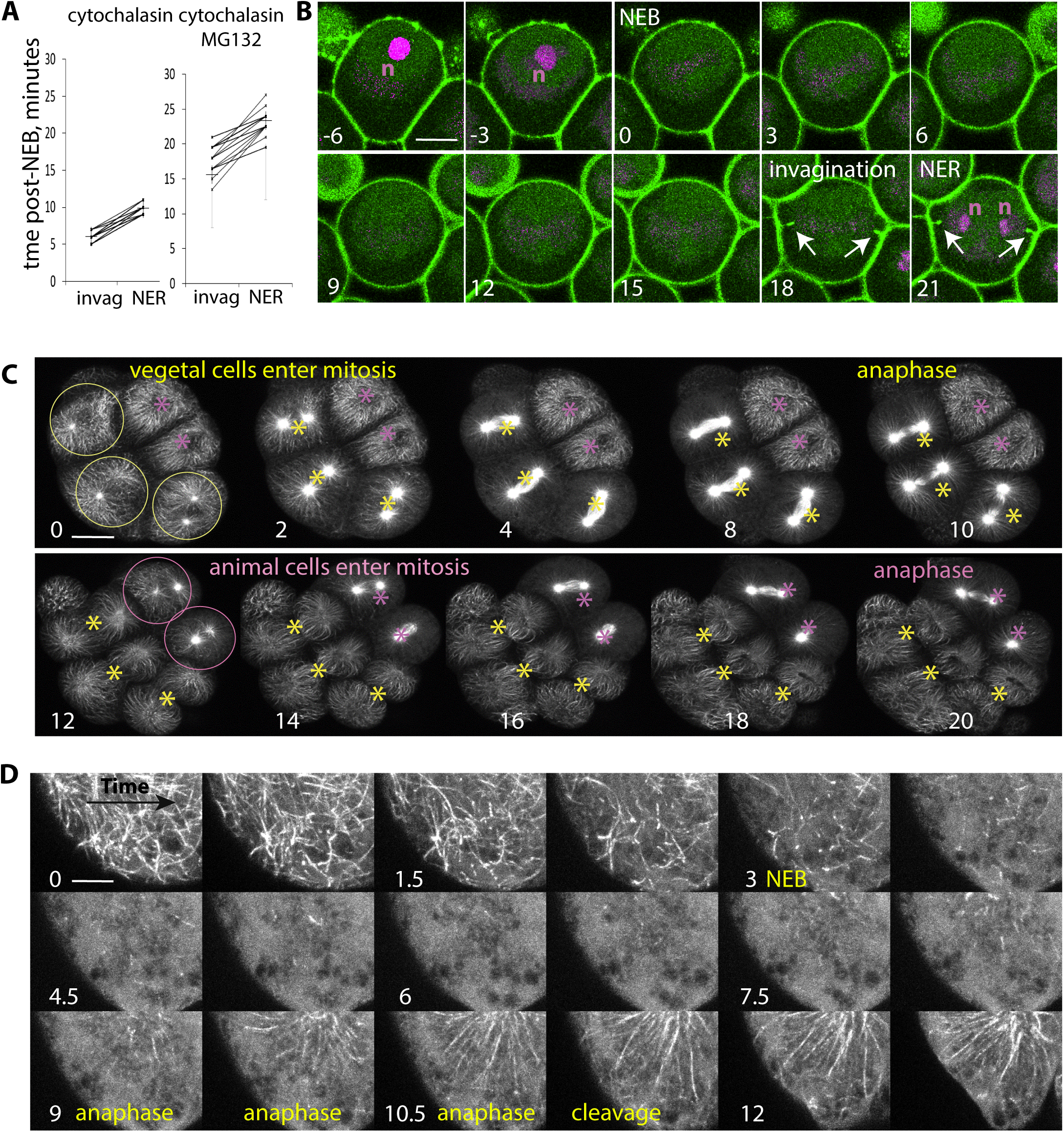
Cortical pulling and microtubules over the cell cycle in 16-cell stage embryos. (Related to Figure 5) A) Time in minutes from NEB to invagination onset or from NEB to NER (nuclear envelope reformation) for cells treated with cytochalasin (left, *n* = 26 cells) or with cytochalasin and MG132 (right, *n* = 72 cells). Lines connect the two measurements for the same cell. B) Images from a timelapse video following one cell in an embryo treated with cytochalasin and the APC/C inhibitor MG132. Plasma membrane is green (PH domain) and nuclei (n) are magenta (NLS). Numbers indicate time in minutes with respect to NEB. Arrows show the emergence of invaginations. NEB: nuclear envelope breakdown; NER: nuclear envelope reformation. Scale bar = 25 microns. C) Images from a timelapse video of a 16-cell stage embryo (anterior view) expressing the microtubule binding protein Map7-GFP (in white). Five cells are visible, 2 from the animal side (pink asterisks) and 3 from the vegetal side (yellow asterisks). First the vegetal side enters mitosis (3 yellow circles, left on top panel), followed by the animal side (2 pink circles, left on bottom panel). Numbers indicate time in minutes. Scale bar = 30 microns. D) Extended display of the timelapse series from Fig. 5 E, showing cyclic disappearance and reappearance of microtubules (labelled with Ensconsin-GFP, white) at the cell surface. Time in minutes (white) and cell cycle events (yellow) are indicated. Scale bar = 8 microns.

## Movie legends

Movie 1: Cortical pulling assay in B5.2 cells. Timelapse fluorescent confocal microscopy of a pair of B5.2 cells treated with cytochalasin, expressing PH-dTomato to label plasma membrane (green) and H2B-GFP to mark chromosomes (magenta). Time interval, 20 seconds.

Movie 2: 3D visualization of midline localization of LGN. A Z-scan of a pair of B5.2 cells fixed and labelled for LGN (green), DNA (blue), and the CAB (anti-Par3, magenta) in anaphase was segmented, reconstructed and pivoted to display the midline patch of LGN with respect to the apical CAB.

Movie 3: Spindle behavior in control embryo. Timelapse fluorescent confocal microscopy (top) and brightfield (bottom) images of the pair of B5.2 cells in a 16-cell *Phallusia* embryo expressing centrosome marker EB3-Venus (green) and H2B-RFP (magenta). Time interval, 60 seconds.

Movie 4: Spindle behavior in LGN-Cter expressing embryo. Timelapse fluorescent confocal microscopy (top) and brightfield (bottom) images of the pair of B5.2 cells in a 16-cell *Phallusia* embryo expressing centrosome marker EB3-Venus (green) and LGN-Cter dominant negative construct. Time interval, 60 seconds.

Movie 5: Cycling of astral microtubules upon entry to and exit from mitosis. Timelapse fluorescent confocal microscopy showing 1 cell from a 16-cell stage *Phallusia* embryo expressing Map7-GFP to label microtubules (green) and H2B-RFP to label chromosomes (red). Time interval, 60 seconds.

